# Non-invasive neuromodulation of cerebello-hippocampal volume-behavior relationships

**DOI:** 10.1101/2024.03.29.587400

**Authors:** Thamires N. C. Magalhães, Ted Maldonado, T. Bryan Jackson, Tracey H. Hicks, Ivan A. Herrejon, Thiago J. R. Rezende, Abigail C. Symm, Jessica A. Bernard

## Abstract

The study here explores the link between transcranial direct current stimulation (tDCS) and brain-behavior relationships. We propose that tDCS may indirectly influence the complex relationships between brain volume and behavior. We focused on the dynamics between the hippocampus (HPC) and cerebellum (CB) in cognitive processes, a relationship with significant implications for understanding memory and motor skills. Seventy-four young adults (mean age: 22±0.42 years, mean education: 14.7±0.25 years) were randomly assigned to receive either anodal, cathodal, or sham stimulation. Following stimulation, participants completed computerized tasks assessing working memory and sequence learning in a magnetic resonance imaging (MRI) environment. We investigated the statistical interaction between CB and HPC volumes. Our findings showed that individuals with larger cerebellar volumes had shorter reaction times (RT) on a high-load working memory task in the sham stimulation group. In contrast, the anodal stimulation group exhibited faster RTs during the low-load working memory condition. These RT differences were associated with the cortical volumetric interaction between CB-HPC. Literature suggests that anodal stimulation down-regulates the CB and here, those with larger volumes perform more quickly, suggesting the potential need for additional cognitive resources to compensate for cerebellar downregulation. This new insight suggests that tDCS can aid in revealing structure-function relationships, due to greater performance variability, especially in young adults. It may also reveal new targets of interest in the study of aging or in diseases where there is also greater behavioral variability.

## 1. Introduction

Recent research has shed light on the cerebellum’s (CB) involvement in both motor and cognitive functions (Strick et al. 2009). Investigations have revealed a functional interplay between the CB and regions of the cerebral cortex responsible for cognitive processing (Schmahmann 2019a; Schmahmann 1996b). Additionally, there is evidence of connections between the CB and the hippocampus (HPC), extending our understanding of cerebellar connections to include subcortical regions as well (Schmahmann 2019a; Watson et al. 2019). Some findings, particularly those concerning links between the CB and HPC have emerged, primarily from animal studies (Watson et al. 2019; Zeidler et al. 2020; Wikgren et al. 2010; Liu et al. 2012). For example, electrical stimulation applied to the CB resulted in the generation of potentials in the HPC in work using rodent models (Zeidler et al. 2020). Animal studies have shown that inhibiting the cerebellar cortex has a significant but regulated impact on the cerebral cortex and subcortical regions, including the HPC (Choe et al. 2018). This effect is due to the multisynaptic connections between the CB and other brain regions (Watson et al. 2019; Bohne et al. 2019; Krook-Magnuson et al. 2014). While evidence suggests CB-HPC connections in rodents (Zeidler et al. 2020; Wikgren et al. 2010), the relationship in humans remains poorly understood.

The interplay between CB and HPC in cognitive processes is of great potential interest, particularly for memory and motor skills. The HPC is known to integrate various information sources, including self-movement cues (McNaughton et al. 2006; van Strien et al. 2009). This integration may rely on indirect input from the CB (Zeidler et al. 2020; Hitier et al. 2014; Rondi-Reig et al. 2014). Recent research using optogenetic functional magnetic resonance imaging (fMRI) has revealed the functional activation of specific forebrain and midbrain regions when pausing Purkinje neurons in the cerebellar cortex’s forelimb region (Choe et al. 2018). Surprisingly, aside from the expected effects on motor-related brain areas, impacts were observed in non-motor regions like the HPC and the anterior cingulate cortex (Choe et al. 2018).

In the context of memory, where the HPC plays a central role (Eichenbaum 2017), it is crucial to also recognize that the CB wields significant influence, particularly in the realms of motor learning and procedural memory (Ding et al. 2012; Leggio et al. 1999; Bernard and Seidler 2014). Any alteration, whether stemming from cerebellar damage or genetic alterations, can profoundly affect the dynamic interplay and functions associated with the HPC (Joyal et al. 1996; Hilber et al. 1998; Rochefort et al. 2011). Recent studies underscore the critical role of functional connectivity between the CB and HPC, especially in tasks related to spatial and temporal processing (Arrigo et al. 2014; Iglói et al. 2015). From a clinical standpoint, these findings suggest that evaluating and addressing structural or network disruptions in the CB-HPC connection may be pivotal in understanding and managing conditions related to memory, motor learning, and procedural skills, such as Alzheimer’s disease (Ding et al. 2012; Fastenrath et al. 2022; Jacobs et al. 2018; Ohlhauser et al. 2019). Moreover, interventions targeting the functional connectivity between the CB and HPC may show promise in developing therapeutic strategies for conditions characterized by deficits in spatial and temporal processing (Yu and Krook-Magnuson 2015). When the CB functions optimally, it efficiently enhances cortical resources, contributing to sustained task performance (Bernard et al. 2020a). Expanding on this concept, it is reasonable to assume that any reduction in cerebellar function may require the recruitment of additional cortical resources to complete a given task.

Recent advancements in noninvasive neuromodulation techniques have opened the door to precise, temporary alterations in brain activity in proscribed regions, allowing for the study of their impact on behavior (Oldrati and Schutter 2018; Ferruci and Priori 2014). Among these techniques, transcranial direct current stimulation (tDCS) stands out as a tool that can modulate cortical excitability by directing the flow of electrical current, potentially facilitating, or inhibiting specific behaviors (Bindman et al. 1964; Antal et al. 2004; Nitsche et al. 2008; Maldonado et al. 2021a). tDCS facilitates the investigation of non-motor functions within the CB, such as its interplay with the HPC (Bohotin et al. 2003). This is expected to lead to heightened reliance on cortical resources, fostering stronger correlations between CB-HPC behavior and volumes, approaching this connection, especially in the context of young individuals, holds the promise of advancing our understanding of various neurological conditions and the aging process.

Recent studies combining tDCS with fMRI have gone a step further by suggesting that stimulating specific areas of the cortical surface can trigger broader changes in the state of interconnected brain regions (Bernard et al. 2020a; Maldonado et al. 2021a; Maldonado et al. 2021b). An illustrative example comes from Hampstead et al. (2014), where parietal-frontal tDCS not only altered activity in the targeted cortical regions but also had observable effects on distant brain areas like the HPC and caudate nucleus (Hampstead et al. 2014). This suggests that tDCS might affect brain function both locally and through network connections, influencing cortical activation patterns, this could offer a way to modulate brain function in a more widespread manner, potentially affecting regions like the HPC (Maldonado et al. 2021a; Hampstead et al. 2014).

Our study aimed to explore the connections between cerebellar volume, HPC, and behavioral performance in young adults. We evaluated their performance in explicit motor sequence and working memory learning tasks. To do this, we paired cerebellar tDCS with structural brain imaging. By using tDCS, a known behavior modulator, we could examine how brain volume interacts with behavior, particularly considering the performance changes often seen after tDCS. The relationship between brain volumes and task performance following tDCS is likely intricate and still little known. Changes in brain structure might underlie enhancements or declines in task performance. We hypothesized that larger volumes of gray matter in both the cerebellum and hippocampus could be linked to improved cognitive function in tasks involving these regions. In our study, participants received one of three types of stimulation (anodal, cathodal, or sham), each targeting the right cerebellar hemisphere. They then completed explicit motor sequence learning and verbal working memory tasks. Our primary goal was to determine whether and to what degree regional brain volume correlated with behavior under simulated and stimulated conditions. Our approach, combining tDCS with volumetric analysis, provides insights into potential relationships and interactions between brain volume and behavior.

## 2. Methods

### 2.1. Participants

Seventy-four young adults (35 males, age: 22±0.42 years, education: 14.7±0.25 years) recruited at Texas A&M University enrolled in this study. Exclusion criteria included left-handedness, history of neurological or mood disorders, skin conditions, and history of concussion. Participants were randomly assigned to either the anodal, cathodal, or sham stimulation condition.

All participants completed a consent form before we initiated any testing procedures. Following the completion of the consent form, participants completed a basic demographic survey. Stimulation was completed followed by the completion of computerized Sternberg (Sternberg 1966) and sequence learning (Kwak et al. 2012) tasks in the magnetic resonance imaging (MRI) environment while brain imaging data were collected. The behavioral data were all collected within 80 minutes of the completion of the stimulation protocol. Tasks were administered in a pre-determined random order. The entire experiment took approximately two hours to complete. All procedures completed by participants were approved by the Texas A&M University Institutional Review Board and conducted according to the principles expressed in the Declaration of Helsinki.

Notably, the analyses presented here are based on a sample of previously collected participants, and aspects of this data have been published already (Maldonado et al. 2021a). In the interest of clarity concerning methods reporting, the methods section below parallels that from our prior work about the behavioral, imaging, and tDCS data collection parameters. The novel data analyses distinguish this work from our prior publication focusing on functional brain activation patterns (Maldonado et al. 2021a).

### 2.2. tDCS Procedures

Participants were blind to the stimulation types. Cathodal, anodal, or sham stimulation was administered using a Soterix 1×1 tES system. Each electrode was placed in a saline-soaked sponge (6 mL per side), with the stimulation electrode placed two cm below and four cm lateral of the inion over the right cerebellum, and the return electrode placed on the right deltoid (Ferruci and Cortese 2015).

To ensure a proper connection with the scalp, an initial 1.0 mA current was set for 30 seconds. If contact quality was below 40%, adjustments, such as moving hair to increase the electrode’s contact with the scalp, were made and contact quality was rechecked. Following a successful re-check, participants completed a 20-minute stimulation session at 2 mA (Ferruci and Cortese 2015; Grimaldi et al. 2016a; Grimaldi et al. 2014b). During the stimulation conditions, maximum stimulation intensity was reached in 30 seconds and maintained for 20 minutes and then would return to 0 mA. During the sham condition, maximum stimulation intensity would be reached, but would then immediately return to 0 mA. There was no additional stimulation during the 20-minute session.

Following anodal, cathodal, or sham stimulation, participants completed a motor learning (explicit sequence learning) or verbal working memory (Sternberg) task in the scanner.

### 2.3. Behavioral Tasks

Participants completed both motor (sequence learning) and non-motor (Sternberg verbal working memory) tasks to better understand how the availability of cerebellar processing resources (as operationalized here by volume) impacts the behavior. Task administration started about 20 minutes after stimulation and the tasks took approximately 35 minutes to complete. This was within the 90-minute window in which the stimulation was thought to be most effective (Nitsche and Paulus 2001). Critically, however, task order was counterbalanced across participants to mitigate any potential impacts of time after stimulation on task performance. All tasks were administered via computer using PsychoPy (Pierce 2007a; Pierce 2019b).

#### 2.3.1. Sequence Learning task

Participants (anodal n=25, cathodal n=24, and sham n=22) were shown four empty rectangles and instructed to indicate the location of the rectangle that was filled as quickly as possible via a button press. Though the stimuli were presented for 200ms, the participant had 800ms to respond before the next stimulus appeared. Random blocks (R) had 18 trials and sequence (S) blocks had 36 trials. During sequence trials, participants had to learn a six-element sequence (1-3-2-3-4-2), which was repeated six times within a block. The order of the task was as follows: R-S-S-S-R-R-S-S-S-R-R-S-S-S-R. For analysis here, the first three sequence blocks were considered early learning, the central sequence blocks were middle learning, and the last sequence blocks were considered late learning (Ballard et al. 2019). Early learning is marked by more cognitively focused activities that necessitate active thinking and working memory (Imamizu et al. 2000; Anguera et al. 2012a; Doyon et al. 1997). As the skill becomes automatic via repetition and practice, the late learning phase becomes increasingly motor-focused. Notably, while the task used here was completed on a shorter timescale, prior work from our group and others has shown that in these laboratory-based tasks, learning typically occurs very quickly (Anguera et al. 2010b; Ballard et al. 2019). Dependent variables used to estimate learning were mean reaction time for correct trials and average total accuracy.

#### 2.3.2. Sternberg task

At the beginning of a trial, participants (anodal n=23, cathodal n=25, and sham n=24) were given six seconds to remember a string of either one, five, or seven capitalized letters, which represent low, medium, and high load, respectively. Following the presentation of the study letters, participants were shown individual lower-case letters and told to indicate whether the letter was one of the study letters shown at the beginning of the trial, via button press. Each letter was displayed for 1200ms, separated by a fixation cross that lasted 800ms. Each participant completed three runs of this task. Within each run, a participant completed three blocks of 25 trials each, for a total of 225 trials. Dependent variables were average reaction time for correct trials and accuracy.

### 2.4. Imaging acquisition

Scanning protocols were adapted from the multiband sequences developed by the Human Connectome Project (HCP) (Ballard et al. 2019; Harms et al. 2018) and the Center for Magnetic Resonance Research at the University of Minnesota to facilitate future data sharing and reproducibility. Data used here are from a larger study wherein participants underwent a fMRI and structural imaging acquisition. The behavioral data used here come from those functional scans. Analyses of the fMRI data have already been published, including an analysis of the behavioral effects of tDCS [34]. Here, our focus is on the structure-behavior relationships. For structural MRI, we collected a high-resolution T1-weighted 3D magnetization prepared rapid gradient multi-echo (MPRAGE) scan (repetition time (TR) = 2400 ms; acquisition time = 7 minutes; voxel size = 0.8 mm^3^).

#### 2.4.1. FreeSurfer Analyses

We used FreeSurfer v.6.0.1, available at https://surfer.nmr.mgh.harvard.edu, to assess the cortical thickness and volume of the CB and HPC regions. Additionally, we investigated measurements of cerebellar white matter (WM) generated by FreeSurfer after image processing. To conduct the analysis, we processed all high-resolution T1-weighted images through the default FreeSurfer processing stages, which included non-linear registration (warping) from the original space to standard MNI space, cortical and subcortical segmentations, and cortical thickness measurements. To smooth the surface, we applied a Gaussian filter with a 10-mm full width at half maximum (Thompson et al. 1997). For this study, we focused on the following segmented regions (bilateral): hippocampus, cerebellum, and cerebellar white matter (WM).

All T1w images were corrected for intensity non-uniformity (INU) with N4BiasFieldCorrection (Tustison et al. 2010), distributed with ANTs 2.3.3 ((Avants et al. 2008), RRID: SCR_004757). The T1w-reference was then skull-stripped with a Nipype implementation of the antsBrainExtraction.sh workflow (from ANTs), using OASIS30ANTs as the target template. Brain tissue segmentation of cerebrospinal fluid (CSF), white matter (WM), and gray matter (GM) was performed on the brain-extracted T1w using fast (FSL 6.0.5.1:57b01774, RRID: SCR_002823, (Zhang et al. 2001)). A T1w-reference map was computed after registration of 2 T1w images (after INU-correction) using mri_robust_template (FreeSurfer 6.0.1, (Reuter et al. 2010)). Brain surfaces were reconstructed using recon-all (FreeSurfer 6.0.1, RRID: SCR_001847, (Fischl et al. 2017)), and the brain mask estimated previously was refined with a custom variation of the method to reconcile ANTs-derived and FreeSurfer-derived segmentations of the cortical gray-matter of Mindboggle (RRID: SCR_002438, (Klein et al. 2017)).

Volume-based spatial normalization to two standard spaces (MNI152NLin6Asym, MNI152NLin2009cAsym) was performed through nonlinear registration with antsRegistration (ANTs 2.3.3), using brain-extracted versions of both T1w reference and the T1w template. The following templates were selected for spatial normalization: FSL’s MNI ICBM 152 non-linear 6th Generation Asymmetric Average Brain Stereotaxic Registration Model [(Evans et al. 2012), RRID: SCR_002823; TemplateFlow ID: MNI152NLin6Asym], ICBM 152 Nonlinear Asymmetrical template version 2009c [(Fonov et al. 2011), RRID:SCR_008796; TemplateFlow ID: MNI152NLin2009cAsym].

### 2.5. Statistical Analysis

For statistical analysis, we used IBM’s SPSS software (version 29, SPSS Inc., Chicago, IL, USA). Initially, we performed a MANCOVA analysis, covarying for age, sex, and estimated total intracranial volume (eTIV), to compare the cortical thickness measures of the cerebellum and hippocampus regions across the stimulation groups. Notably, we did not predict any group differences but investigated this to determine the degree of uniformity in volumetric measures across groups.

Subsequently, we conducted partial correlations between task measures and cortical thickness measures while controlling for eTIV. Moreover, we examined the statistical interaction between hippocampal and cerebellar volumes and task performance following tDCS, inspired by the cerebellar-hippocampal circuit outlined in the animal literature (Zeidler et al. 2020). Studying the interaction between volumetric cortical CB-HPC has the potential to deepen our understanding of memory, learning, and sensorimotor integration, as well as to provide insights into neurological and psychiatric disorders. However, it’s crucial to clarify that our study specifically examined the statistical interactions of these regional volumes. We included this analysis because these brain regions are integral parts of a larger network that underpins a range of cognitive and behavioral functions. Our goal in examining these statistical interactions was to reveal the collaborative nature of these regions in information processing and behavioral contributions. We established cross-lateralized interactions (between the cortical volume of the regions): *Right cerebellum*Left hippocampus (CB-HPC R*L) and Left cerebellum*Right Hippocampus (CB-HPC L*R)*.

Following the completion of the initial partial correlation analysis, our subsequent analyses involved performing a hierarchical multiple regression to explore the predictive relationship between post-tDCS behavioral measures and brain volume metrics. Before this analysis, we conducted preliminary assessments to ensure that the assumptions of normality, linearity, multicollinearity, and homoscedasticity were satisfied. After confirming these assumptions, we proceeded with linear regression models for brain regions that displayed significant correlations with the sequence learning and Sternberg task measurements. In this modeling, we designated task measures as the dependent variable and cortical segmentation measures as the independent variable. This methodological strategy enabled us to enhance our comprehension of how the tDCS stimulation used affects interactive effects. To address the issue of multiple comparisons, we implemented a Bonferroni correction, as mentioned in each result of the analysis obtained and when the correction was necessary.

## 3. Results

Our group has previously published task performance differences between stimulation groups. According to Maldonado et al. 2021, regarding the sequence task, reaction times (RT) were found to be significantly higher in the cathodal group compared to the anodal and sham groups. Furthermore, the learning phase had a significant impact, with RTs differing significantly between the early, intermediate, late, and random learning measures (Maldonado et al. 2021a). As for the Sternberg task, the effects were primarily seen with the load conditions with RT varying between different load levels, demonstrating the success of the load manipulation. Additionally, there was a notable effect of stimulation, as RTs in both the anodal and cathodal groups differed from those in the sham group (Maldonado et al. 2021a).

To examine the potential variation in volumes across groups, we conducted a MANCOVA analysis (Wilks’ Lambda = 0.02) as outlined in **Table 1**. Interestingly, the left hippocampal volume was found to be significantly lower in both the cathodal and anodal groups when compared to the sham group. Additionally, disparities in left cerebellar WM volume were observed in the cathodal group in contrast to the sham group, indicating a smaller volume in the cathodal group. It is essential to note, however, that these differences likely stem from the inherent variability within our study population, as group assignments were made randomly. Notably, there were no discernible variations among the groups concerning age and education level. Furthermore, no significant differences were detected regarding the interactions between cerebellar and hippocampal volumes among the groups.

**Table 1.**
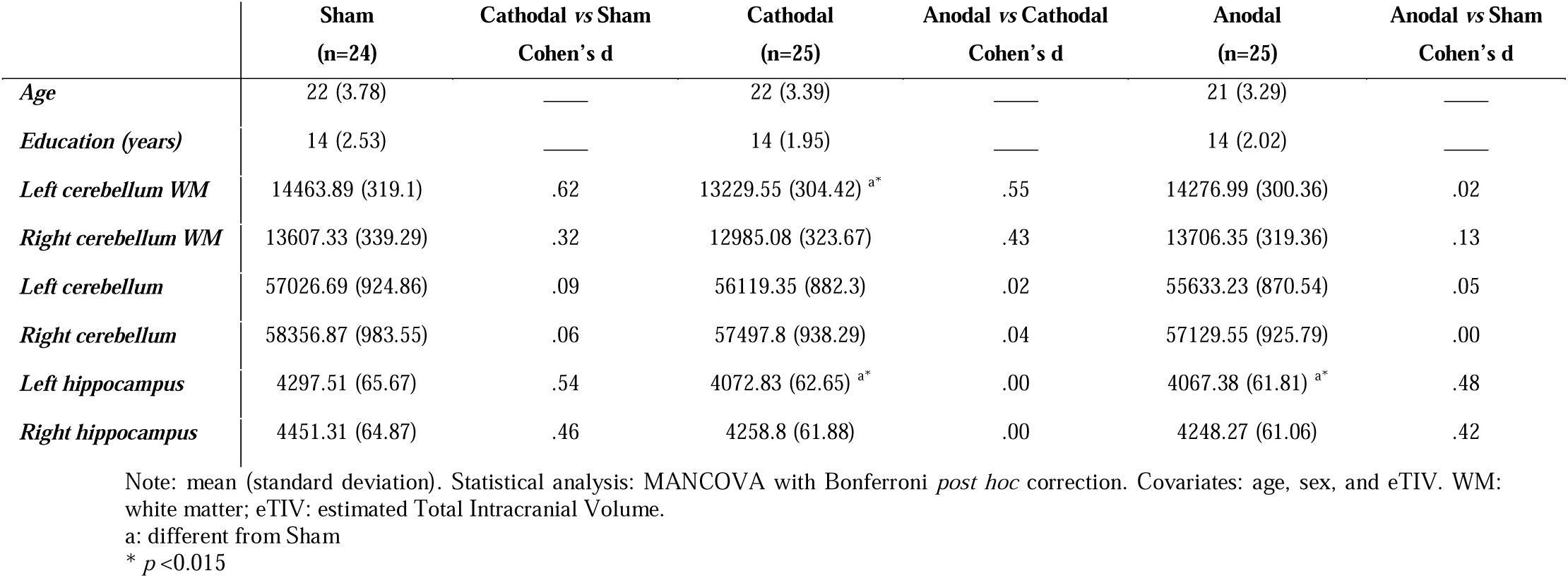
Comparisons of cortical segmentation and demographic data across groups.

### 3.1. Correlations between tasks and cortical segmentation

#### 3.1.1. Sequence task

Though there were patterns indicating some relationships between brain structure and behavioral performance on the sequence learning task, these results were not significant following Bonferroni correction for multiple comparisons (p<0.015). For the complete listing of results please see **Table 3**.

**Table 3.** Correlations analysis between stimulation groups and cortical volume.

| Tasks | Group | Left-Cerebellum | Right-Cerebellum | Left-Hippocampus | Right-Hippocampus | CB-HPC (L*R) | CB-HPC (R*L) |
| --- | --- | --- | --- | --- | --- | --- | --- |
| *mean acc Sequence* | Sham | .142; $p=0.53$ | .320; $p=0.15$ | .085; $p=0.71$ | -.133; $p=0.56$ | 0.280; $p=0.22$ | .002; $p=0.99$ |
| | Cathodal | .200; $p=0.37$ | .111; $p=0.62$ | -.460; $p=0.031$ | -.389; $p=0.073$ | -.215; $p=0.33$ | -.070; $p=0.75$ |
| | Anodal | .290; $p=0.18$ | .268; $p=0.21$ | -.122; $p=0.57$ | -.130; $p=0.55$ | .093; $p=0.67$ | .109; $p=0.62$ |
| *mean rt Sequence* | Sham | -.283; $p=0.21$ | -.423; $p=0.056$ | .125; $p=0.58$ | -.096; $p=0.67$ | -.193; $p=0.4$ | -.262; $p=0.25$ |
| | Cathodal | -.160; $p=0.47$ | .006; $p=0.97$ | -.419; $p=0.052$ | -.277; $p=0.21$ | -.273; $p=0.21$ | -.268; $p=0.22$ |
| | Anodal | -.357; $p=0.094$ | -.336; $p=0.11$ | -.238; $p=0.27$ | -.289; $p=0.18$ | -.332; $p=0.12$ | -.374; $p=0.07$ |
| *mean random rt Sequence* | Sham | -.189; $p=0.41$ | -.327; $p=0.14$ | -.050; $p=0.83$ | -.264; $p=0.24$ | -.257; $p=0.26$ | -.311; $p=0.17$ |
| | Cathodal | -.206; $p=0.35$ | -.027; $p=0.9$ | -.381; $p=0.08$ | -.249; $p=0.26$ | -.294; $p=0.18$ | -.312; $p=0.15$ |
| | Anodal | -.425; $p=0.026$ | -.466; $p=0.026$ | -.168; $p=0.44$ | -.252; $p=0.24$ | -.370; $p=0.08$ | -.418; $p=0.047$ |
| *mean late learning rt Sequence* | Sham | -.264; $p=0.24$ | -.382; $p=0.08$ | .077; $p=0.73$ | -.137; $p=0.55$ | -.192; $p=0.4$ | -.271; $p=0.23$ |
| | Cathodal | -.187; $p=0.4$ | -.035; $p=0.87$ | -.446; $p=0.038$ | -.198; $p=0.37$ | -.314; $p=0.15$ | -.237; $p=0.28$ |
| | Anodal | -.190; $p=0.38$ | -.156; $p=0.47$ | -.320; $p=0.13$ | -.404; $p=0.056$ | -.273; $p=0.2$ | -.345; $p=0.1$ |
| *mean high rt Sternberg* | Sham | **<u>-.533; $p=0.013$</u>** | -.489; $p=0.025$ | .120; $p=0.6$ | .014; $p=0.95$ | -.226; $p=0.32$ | -.347; $p=0.12$ |
| | Cathodal | .000; $p=0.99$ | .145; $p=0.51$ | .328; $p=0.13$ | .417; $p=0.053$ | .324; $p=0.14$ | .251; $p=0.25$ |
| | Anodal | .288; $p=0.18$ | .252; $p=0.25$ | .096; $p=0.66$ | .038; $p=0.86$ | .221; $p=0.31$ | .195; $p=0.37$ |
| *mean low rt Sternberg* | Sham | -.240; $p=0.29$ | -.350; $p=0.12$ | .102; $p=0.65$ | -.174; $p=0.45$ | -.155; $p=0.5$ | -.276; $p=0.22$ |
| | Cathodal | -.059; $p=0.79$ | .113; $p=0.61$ | -.162; $p=0.47$ | -.136; $p=0.54$ | -.069; $p=0.76$ | -.152; $p=0.49$ |
| | Anodal | -.415; $p=0.049$ | -.352; $p=0.099$ | -.181; $p=0.41$ | -.407; $p=0.054$ | -.325; $p=0.13$ | **<u>-.516; $p=0.012$</u>** |

Note: Statistical analysis: Partial correlations. Covariates: eTIV. Acc: accuracy; RT: reaction time; eTIV: estimated Total Intracranial Volume. Bold underlined values: results that remained significant after corrections for multiple comparisons ( $p < 0.015$ ).

#### 3.1.2. Sternberg working memory task

Following correction for multiple comparisons (p < 0.015), we observed negative correlations between the left cerebellar cortical volume and mean RT during high-load blocks in the sham group, with correlation coefficients of -.533 (p = 0.013), as illustrated in **Figure 1**. In the anodal stimulation group, we also detected correlations between mean RT during low-load blocks and the CB-HPC *R*L* interaction (r = -.516, p = 0.012). For a comprehensive overview of the results, please refer to **Table 3**.

**Figure 1.**
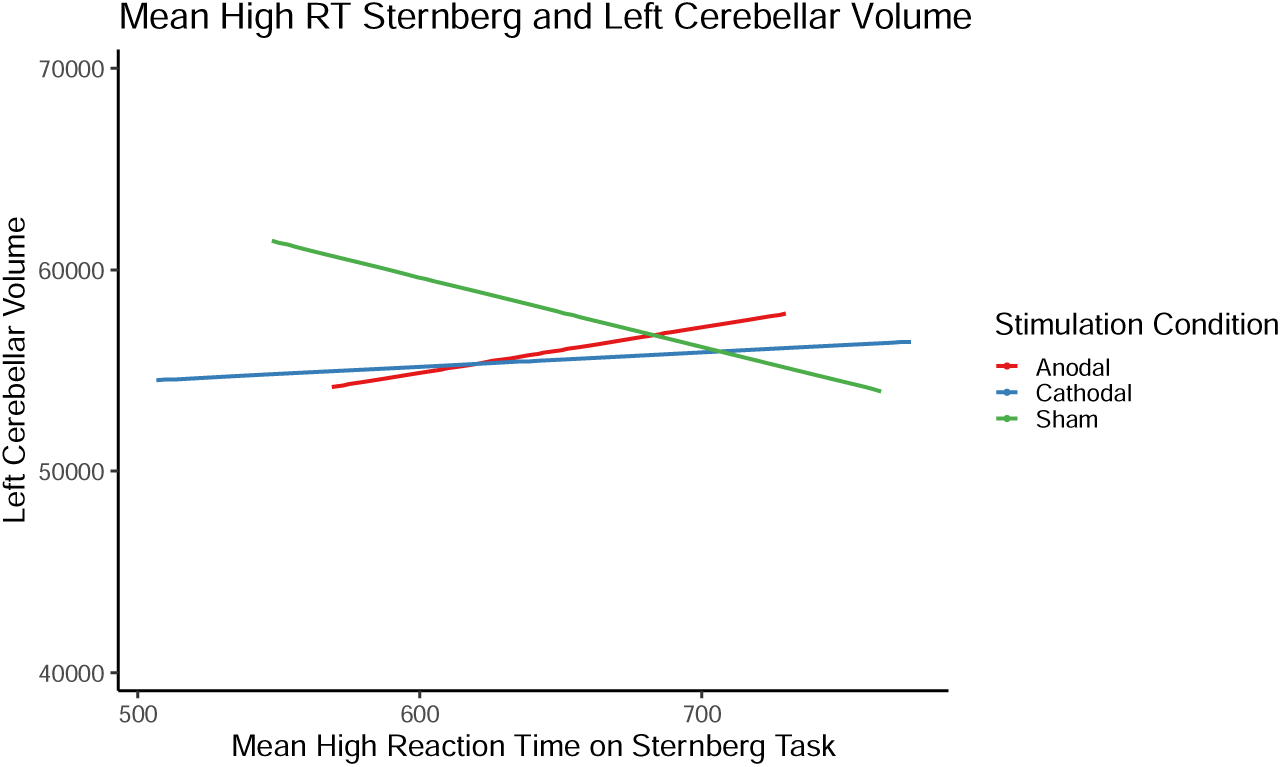
Illustrated the correlation analysis between mean scores of high reaction time (RT) in the Sternberg test and the left cerebellar volume across three different stimulation groups (Sham, Cathodal, and Anodal). Notably, the green line representing the Sham group yielded a significant result.

### 3.2. Linear regression analysis

After examining the correlations and pinpointing variables with significant relationships, specific to the Sternberg task, we conducted stepwise linear regression analyses. This aimed to delve deeper into understanding how brain structure (volume) is influenced after stimulation, offering a more comprehensive analysis of the brain-behavior relationship.

#### Sham group

In the initial block, the inclusion of eTIV explained less than 1% of the variance in mean RT during the high load condition. Upon introducing cerebellar cortical volume (right and left hemisphere) and hippocampal volume (right and left) in the second block, the model’s total variance increased to 36%, though the model itself was not statistically significant (F(5, 16) = 1.81, p < .16). In this model, no cortical measures achieved statistical significance, and, while the volume measures accounted for a greater proportion of the variance, the overall final model was not statistically significant.

#### Anodal group

In the initial block, eTIV was included and again accounted for less than 1% of the variance in mean RT during low-load blocks. Upon introducing CB-HPC *L*R* and *R*L* volume measures (representing interactions, as detailed in Section *2.5*, *Statistical Analysis*) in the second block, the model explained a total variance of 39% (F (3, 20) = 4.29, p = .017). The incorporation of CB-HPC *L*R* and *R*L* interactions measures contributed nearly 38% additional variance in the mean low RT measure, even after controlling for eTIV, as evidenced by R squared change = .3 and F change (2, 391 = 6.43, p < .02). In the final model, only one measure, the CB-HPC *R*L* (beta = –2.06, p < .001), was found to be statistically significant, as illustrated in **Figure 2**.

**Figure 2.**
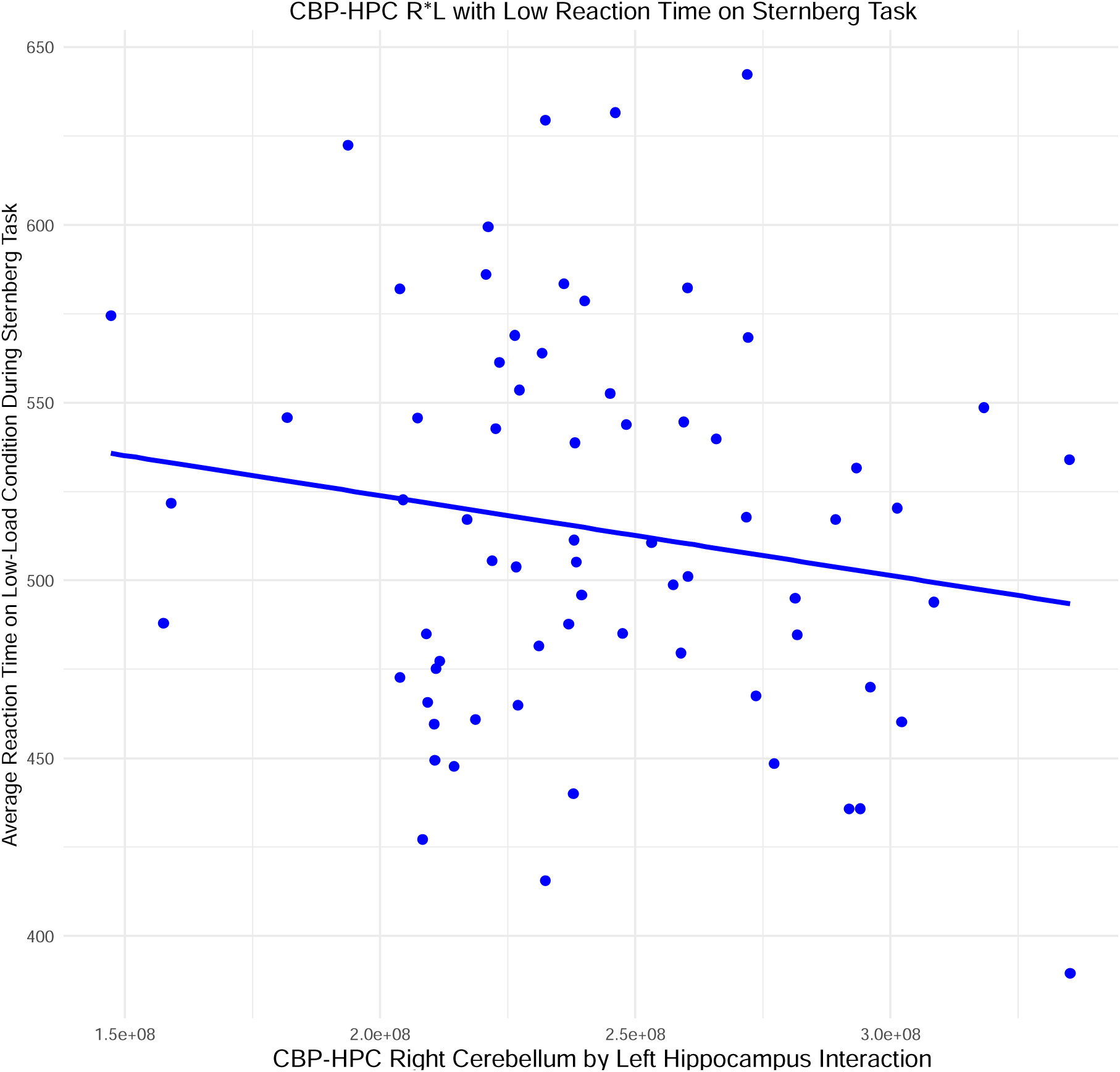
The anodal stimulation group exhibited a cortical volumetric interaction between the right cerebellum and left hippocampus (CB-HPC *R*L*) during the mean reaction time (RT) in low-load blocks of the Sternberg test.

## 4. Discussion

Our investigation provides valuable insights into the dynamic relationships between the cerebellar and hippocampal structure and behavioral performance, especially within the context of cerebellar stimulation. Within the sham group, we noted a correlation between cerebellar volume and working memory. Individuals with larger cerebellar volumes demonstrated faster RT during high-load conditions, implying that larger cerebellar volumes could enhance cognitive processing efficiency. In contrast, the group that received anodal cerebellar stimulation exhibited a distinct pattern. In this group, lower RT during the low load condition of the Sternberg task was linked to the volumetric interaction between the cerebellum and hippocampus. These findings suggest that the interaction between these brain regions may play a crucial role in enhancing or sustaining task performance in individuals undergoing anodal stimulation.

In their recent study, Maldonado and colleagues found that anodal cerebellar stimulation during the Sternberg task led to increased cortical activation (Maldonado et al. 2021a). This could be a compensatory response to the reduced output from the CB and its associated processing. Their results suggest that anodal stimulation may impair cerebellar processing needed for cognitive tasks, resulting in higher activation in other brain regions involved in working memory (Maldonado et al. 2021a). In our study, we observed significant CB-HPC interactions in the anodal stimulation group during the Sternberg task. This supports the idea that anodal stimulation can decrease cerebellar activity, as well as activity in other important regions for cognition (Hampstead et al. 2014; Liebrand et al. 2020; Khan et al. 2020). Our findings highlight the CB’s role in supporting cognitive processes and show that reducing cerebellar output can affect both function and behavior (Emch et al. 2019; Schmahmann et al. 2019; Jonides et al. 1997). Our research contributes to understanding how brain structure, non-invasive stimulation, and cognitive performance are interconnected.

Prior to exploring the potential impact of tDCS on the connections between brain volume and behavior, it is crucial to underscore the significance of our findings concerning the CB-HPC interaction. Our observations within the anodal stimulation group align with the hypothesis proposing that the cerebellar region may enlist other brain areas to access additional cognitive resources, a concept referred to as the scaffolding hypothesis (Maldonado et al. 2019b; Reuter-Lorenz and Park et al. 2014; Filip et al. 2019). According to this hypothesis, when cerebellar function is negatively modulated, supplementary cognitive resources are recruited to maintain or enhance cognitive performance (Filip et al. 2019; Bernard et al. 2013b). The idea of leveraging additional cognitive resources implies that the brain dynamically adjusts to sustain optimal performance, even in the face of temporary alterations to specific neural pathways. This adaptability holds practical implications, such as in developing therapeutic interventions for conditions involving cerebellar dysfunction, like diseases, or in optimizing cognitive performance in healthy individuals.

The interaction between the CB and HPC represents a complex and multifaceted aspect of brain function. While our understanding continues to evolve, several key points underscore the importance of investigating this interaction. Although the cerebellum is conventionally associated with motor control and coordination, recent research has convincingly demonstrated its involvement in non-motor functions and cognitive processes (Strick et al. 2009; Buckner et al. 2013). The hippocampus, known for its role in memory formation and spatial navigation, is also a region commonly affected by neurodegenerative processes (Devanand et al. 2007). Exploring this interaction significantly contributes to our understanding of various aspects of cognitive processing, encompassing tasks related to working memory, attention, and executive functions (Rondi-Reig et al. 2014; Rochefort et al. 2011). Disruptions in the CB-HPC interaction have been implicated in several neurological and psychiatric disorders, shedding light on the underlying mechanisms of conditions such as schizophrenia (Bernard and Mittal et al. 2015) and neurodegenerative diseases (Hoxha et al. 2018; Olivito et al. 2020). As mentioned earlier, the brain’s adaptability, as evidenced by the scaffolding hypothesis, emphasizes the dynamic nature of these interactions and their profound impact on overall brain function.

While numerous studies have focused on the impact of tDCS on cortical excitability and functional alterations, as evidenced in fMRI studies (Choe et al. 2019; Maldonado et al. 2021a; Hampstead et al. 2014; Emch et al. 2019), a noticeable research gap persists in assessing the connections between brain volume and behavioral performance. The advantages of tDCS, including its affordability, safety, portability, and ease of use, have significantly increased its utilization in both research and clinical settings (Benwell et al. 2015; Vergallito et al. 2022). Given the profound potential implications of tDCS on brain function, it is reasonable to hypothesize that it may indirectly influence relationships between brain volume and behavior, perhaps through behavioral variability, and different structures related to disturbances due to variability.

The observed variability in tDCS effects has been attributed to substantial differences in stimulation protocols across studies, encompassing variations in stimulation parameters, target regions, and electrode montage (Vergallito et al. 2022). Beyond these factors, the sequencing of tasks and the temporal gap between the tDCS session and the two tasks conducted in the scanner may introduce a potential variable influencing behavioral outcomes (Vergallito et al. 2022). Despite conscientious attempts to counterbalance the task order and mitigate the potential impact of diminishing stimulation effects, it is still possible that the influence of stimulation was somewhat diminished during the second task within the scanner. Previous studies showed that this did not have a significant impact (Maldonado et al. 2021a). Furthermore, our findings unearthed noteworthy discrepancies in brain volume across distinct stimulation groups. The diverse responses to tDCS may be significantly influenced by individual differences in brain anatomy.

The literature on tDCS does not necessarily address interindividual differences in its effects. The importance of investigating this and its complexities in research design and interpretation is crucial for a more accurate understanding of the effects of tDCS. Interindividual variability in response to tDCS can be attributed to several characteristics, including morphological and genetic factors (Benwell et al. 2015). Genotypic differences can alter the effect of tDCS, influencing the anatomical and neurophysiological states of individuals (Li et al. 2015). For example, carriers of the Met BDNF polymorphism, in an electroencephalogram (EEG) study, showed differences in regional brain volumes and task-specific synchrony, which predicted performance on an error-processing task (Soltész et al. 2014). As for morphological differences, which include volumetric differences between individuals, although it is important when considering stimulation in healthy individuals, which is the case in our study, it becomes much more significant when considering patients with brain injuries. Mahdavi et al. 2018 demonstrated that a decline in gray matter volume, observed in both mild cognitive impairment (MCI) and physiological aging, led to a diminished current density reaching the brain when contrasted with younger participants (Mahdavi et al. 2018a). When considering Alzheimer’s patients, the same authors (Mahdavi et al. 2018b) proposed that structural alterations might modify the regions being stimulated and the location of the maximum current density within the head. Such alterations on this scale have the potential to influence the anticipated behavioral effects of applying tDCS.

Based on these discoveries, it is imperative to illustrate whether the morphological variances influencing the quantity and distribution of the induced electric field can indeed affect the behavioral outcomes of tDCS. Filmer et al. 2019 found that cortical architecture variations predicted the impact of anodal tDCS on behavioral performance. In their study involving 47 healthy young adults, anodal or cathodal tDCS was applied over the left pre-frontal cortex during a decision-making task. Results indicated that individuals with a thicker cortex in specific areas and a thinner cortex in another showed greater disruption of learning with anodal stimulation. Interestingly, interindividual differences did not explain the variability in cathodal tDCS efficacy, implying potential differences in sources of variance for the two stimulation polarities (Filmes et al. 2019).

In summary, our research investigated the volumetric interaction between cerebellar-hippocampal structures and their impact on behavioral performance. We observed that individual volumetric variations played a crucial role in determining distinct RTs, aligning with findings from Filmer and colleagues. Specifically, individuals with larger cerebellar volumes exhibited swifter reaction times during tasks with high cognitive demands, indicating a potential contribution of a larger cerebellum to more efficient cognitive processing.

It is essential to note, however, that further studies are necessary, particularly focusing on young individuals. This emphasis on diverse age groups is crucial for a comprehensive exploration of these relationships and to refine the precision of results, especially concerning their relevance to neuropsychiatric and neurodegenerative diseases. This avenue of research holds promise for advancing our understanding of cognitive processes and their implications for various health conditions.

## 5. Conclusion

Our investigation explores the intricate volumetric interactions between cerebellar and hippocampal structures, behavioral performance, and the influence of anodal cerebellar stimulation. Significant findings support the idea that anodal stimulation modulates cognitive processing by dynamically recruiting additional cognitive resources, aligning with the scaffolding hypothesis. The study contributes to understanding the non-motor functions of the cerebellum and underscores the importance of the CB-HPC interaction in various cognitive processes, with potential implications for neurological and psychiatric disorders.

Furthermore, our research highlights the often-overlooked inter-individual differences in tDCS effects, emphasizing the role of morphology. This understanding is crucial for interpreting tDCS effects accurately, particularly in brain-damaged patients. The study emphasizes the importance of considering morphological differences in the induced electrical field, shedding light on potential impacts on tDCS-induced behavioral changes.

## 6. Funding

This work was supported by the National Institute on Aging, Grant number: R01AG064010.

## 7. Competing Interests

The authors have no relevant financial or non-financial interests to disclose.

## 8. Data Availability

The datasets generated during and/or analyzed during the current study are not publicly but are available from the corresponding author upon reasonable request.

## Notes

### Competing Interest Statement

The authors have declared no competing interest.

